# The CX3CL1 oligomerization is required for efficient CX3CR1-specific cell adherence

**DOI:** 10.1101/865998

**Authors:** Mariano A. Ostuni, Patricia Hermand, Emeline Saindoy, Noëlline Guillou, Julie Guellec, Audrey Coens, Claude Hattab, Elodie Desuzinges-Mandon, Anass Jawhari, Soria Iatmanen-Harbi, Olivier Lequin, Patrick Fuchs, Jean-Jacques Lacapere, Christophe Combadière, Frédéric Pincet, Philippe Deterre

## Abstract

During inflammatory response, blood leukocytes adhere to the endothelium. This process involves numerous adhesion molecules, including a transmembrane chemokine, CX3CL1. We previously found that CX3CL1 clusters in oligomers. How this cluster assembles and whether it has a functional role remain unknown. Using various biochemical and biophysical approaches, we show that CX3CL1 clusters are homo-oligomers with 3 to 7 CX3CL1 molecules. We demonstrate that the transmembrane domain peptide self-associates at a similar level in both cellular and acellular lipid environments while its random counterpart (a scrambled peptide) does not. Hence, oligomerization is mainly driven by the transmembrane domain intrinsic properties. Molecular modeling suggests that transmembrane peptide oligomers are mostly made of monomers linearly assembled side by side. Using a new adherence assay, we demonstrate that, functionally, oligomerization is mandatory for the adhesive potency of CX3CL1. Our results indicate that CX3CL1-dependent cellular adherence in key immune processes can be controlled by disrupting clusters using heterotopic peptides, which, in turn, alter the adhesive function of the membrane CX3CL1 without affecting the function of the CX3CL1 soluble form.

## Introduction

The migration of blood leukocytes to damaged tissues is the first step of the inflammation process and involves a sequence of coordinated interactions between leukocytes and endothelial cells (Ingersoll et al, 2011; Langer & Chavakis, 2009; Luster et al, 2005). The chemotactic cytokines called chemokines that primarily attract leukocytes, are central to the physiological and pathological inflammatory processes (Charo & Ransohoff, 2006; Ortega-Gomez et al, 2013; Ransohoff, 2009). Chemokines trigger leukocyte activation and their firm adhesion to the inflamed endothelium, mainly through integrins (Combadière et al, 2007; Simon et al, 2009; Speyer & Ward, 2011). Two members of the chemokine family are exceptions: CXCL16 and CX3CL1. In addition to their chemokine domain (CD), these two chemokines possess three domains: a mucin-like stalk, a transmembrane (TM) domain, and a cytosolic tail (Bazan et al, 1997; Matloubian et al, 2000). When interacting with their cognate receptors (CXCR6 and CX3CR1, respectively), these chemokines induce cell-cell adhesion (Ludwig & Weber, 2007). CXCL16 and CX3CL1 can also be cleaved by metalloproteinases, such as ADAM10 and ADAM-17 (Garton et al, 2001; Hundhausen et al, 2003; Ludwig et al, 2005), to produce a soluble form with chemotactic functions.

The CX3CL1 chemokine, with its unique CX3CR1 receptor (Imai et al, 1997), is involved in adherence to the endothelium of the inflammatory monocyte population (CD14^hi^ CD16^−^ CX3CR1^+^ CCR2^+^ in humans, Ly6C^hi^ CX3CR1^+^ CCR2^+^ in mice) (Ancuta et al, 2003; Babendreyer et al, 2017; Geissmann et al, 2003; Ludwig & Weber, 2007; Tacke & Randolph, 2006) likely through interaction with platelets (Postea et al, 2012; Schulz et al, 2007). This chemokine is also involved in the recruitment of NK lymphocytes (Guo et al, 2003; Lavergne et al, 2003) and in lymphocyte survival as in allergic diseases (Julia, 2012), as well as in monocytic (Collar et al, 2018; White et al, 2014) and neuronal survival (Cipriani et al, 2011; Kim et al, 2011; Meucci et al, 2000; Mizuno et al, 2003). An additional function of the CX3CR1-CX3CL1 pair is the regulation of the patrolling behavior and the margination of monocytes in blood vessels (Auffray et al, 2007; Hamon et al, 2017) or their adherence to the bone marrow (Jacquelin et al, 2013). The CX3CL1 chemokine is also involved in cytoadhesion of red blood cells infected with the malaria parasite *Plasmodium falciparum* (Hermand et al, 2016). The transmembrane form of CX3CL1, as well as the other transmembrane chemokine CXCL16, is also involved in glial cross-talk (Ludwig & Mentlein, 2008), possibly by a new mechanism called inverse signaling (Hattermann et al, 2016).

We previously showed that two molecular features are involved in the CX3CL1 adhesive function, i.e. the glycosylation of mucin stalk and the intracellular domain. Glycosylation ensures the accessibility of CX3CL1 to the CX3CR1 molecules buried in the membrane of the counter-adhesive cell while the intracellular domain anchors the transmembrane chemokine in its cell membrane (Ostuni et al, 2014). We also showed that the CX3CL1 molecule self-assembles and that this assembly is mainly dependent of its TM domain (Hermand et al, 2008). This leads to a model in which the formation of adhesive patches is controlled by the dynamics of the CX3CR1 receptors that bind the CD domain presented by CX3CL1 bundles. (Ostuni et al, 2014). However, the exact nature of the CX3CL1 assembly is not known, as well as the number of CX3CL1 monomer in each functional bundle. Using a large panel of biophysical, biochemical and cell biology tools, we show here that the CX3CL1 self-assembly is precisely due to the oligomerizing feature of its TM domain and that the oligomerization degree is between 3 and 7. *In silico* molecular modelling suggests that CX3CL1 oligomers are linearly organized. Finally, using the TM domain peptide alone, we were able to specifically modulate the CX3CL1-CX3CR1 dependent cellular adherence, opening the way to new class of inhibitors able to antagonize the function of the CX3CL1 membrane form without affecting the role of the CX3CL1 soluble form.

## Results

### Aggregation degree of the whole CX3CL1

To determine the number of monomers per each CX3CL1 bundle, we first performed electrophoresis of the cell homogenates of L_929_ cells stably expressing CX3CL1 (hitherto denoted L_CX3CL1_) fused to EYFP at its C-terminal (Ostuni et al, 2014). SDS-PAGE showed that the CX3CL1-YFP monomer is around 130 kDa (Fig 1A). Then, using native electrophoresis after solubilization of L_CX3CL_ cell homogenates in dodecylmaltoside (DDM) (Fig 1B), we found an intense band around 480 kDa, i.e. ca. three-fold the molecular weight of the monomer. Another band is also visible around 1100 kDa. Using a milder detergent, digitonin, we found very similar results (Fig EV1). We next used another type of detergent (CALX173ACE from CALIXAR) that was shown to maintain proteins oligomeric state (Agez et al, 2019; Desuzinges Mandon et al, 2017; Igonet et al, 2018) and protein-protein interactions (Rosati et al, 2015). To avoid unspecific staining, the CX3CL1-EYFP was immune-purified before native electrophoresis. We found a major band at 480 kDa and a minor one at 700 kDa, i.e. 5-fold of the monomer molecular weight, while a minor proportion of the protein does not enter the gel (Fig 1C). Taken together our native electrophoresis experiments point out that the native CX3CL1 is certainly not monomeric and contains at least three monomers.

**Fig 1.**
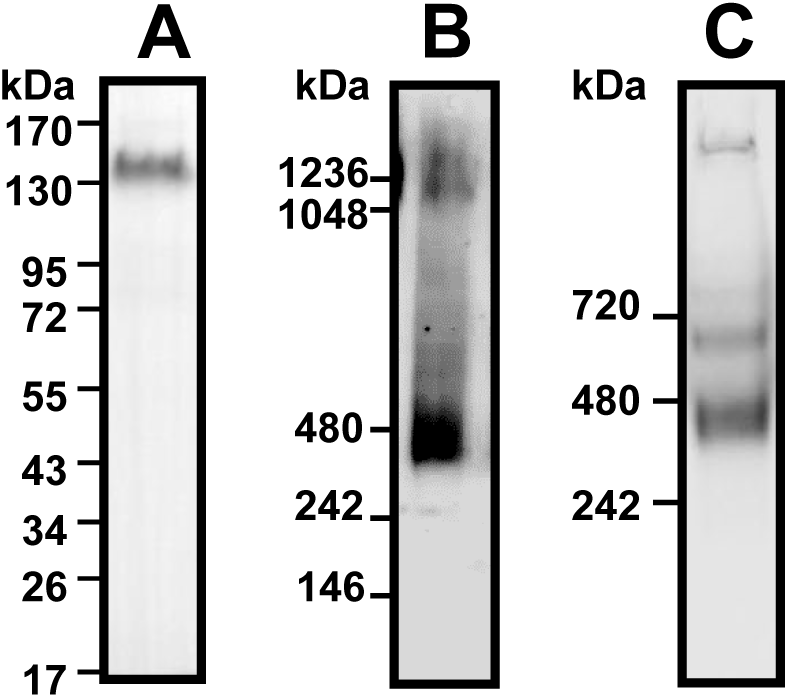
Electrophoresis of the CX3CL1-YFP fusion protein. A.Cell lysate of L_CX3CL1_ cells stably transfected with CX3CL1-EYFP was analyzed by SDS-PAGE and Western blot. B.Cell lysate of L_CX3CL1_ was analyzed by native electrophoresis using Nu-Page gel in the presence of 1% DDM (dodecylmaltoside) and Western blot. C. The affinity-purified CX3CL1 from L_CX3CL1_ membranes was analyzed by native PAGE. CXCL3 was solubilized using CALX173ACE and affinity purified (CXCL1 antibody). Mini-proteon TGX gel electrophoresis (Biorad) were used for Native PAGE.

We next used the single-molecule TIRF microscopy, a reliable technique to investigate the number of subunits in membrane protein complexes (Dietz et al, 2013; Hastie et al, 2013; Hines, 2013; Teramura et al, 2006; Ulbrich & Isacoff, 2007; Zhang et al, 2010). This assay was based on the measurement of spontaneous step-wise decrease of particle fluorescence, assuming that the number of steps is directly related to the number of fluorophores per particles. This technique requires that the fluorescent particles be at low density to ensure they are spatially separated (Ulbrich & Isacoff, 2007). A mixture of L_CX3CL1_ homogenates and 1,2-dioleoyl-*sn*-glycero-3-phosphocholine (DOPC) liposomes was spread on a coverslip and observed by TIRF microscopy (Fig 2A). DOPC was in large excess to ensure spatial separation of the CX3CL1 clusters that appeared as isolated particles. The fluorescence kinetics of more than hundred particles were analyzed. Some particle bleached in one step (Fig 2B left), some in two (Fig 2B center) or more up to 7 steps (Fig 2B right). We noticed that the step amplitude of the particles bleaching in 1 or 2 steps were substantially higher than the amplitude of the fluorescence step of the particles bleaching in 3 steps or more (Fig 2C). This pointed out that in these particles two or more chromophores were bleaching at the same time, indicating that each L_CX3CL1_ particle would contain three EYFP or more. We hypothesized that the mean amplitude of the fluorescence steps of particles bleaching in 4 steps or more (calculated to be 110 a.u.) corresponded to the bleaching of a single dye. So the initial fluorescence intensity of each particle divided by this unit step was a good estimate of the actual number of dyes in each cluster. The resulting distribution (Fig 2D) of the cluster sizes indicated that CX3CL1 forms an oligomer containing 4 ± 2 monomers.

**Fig 2.**
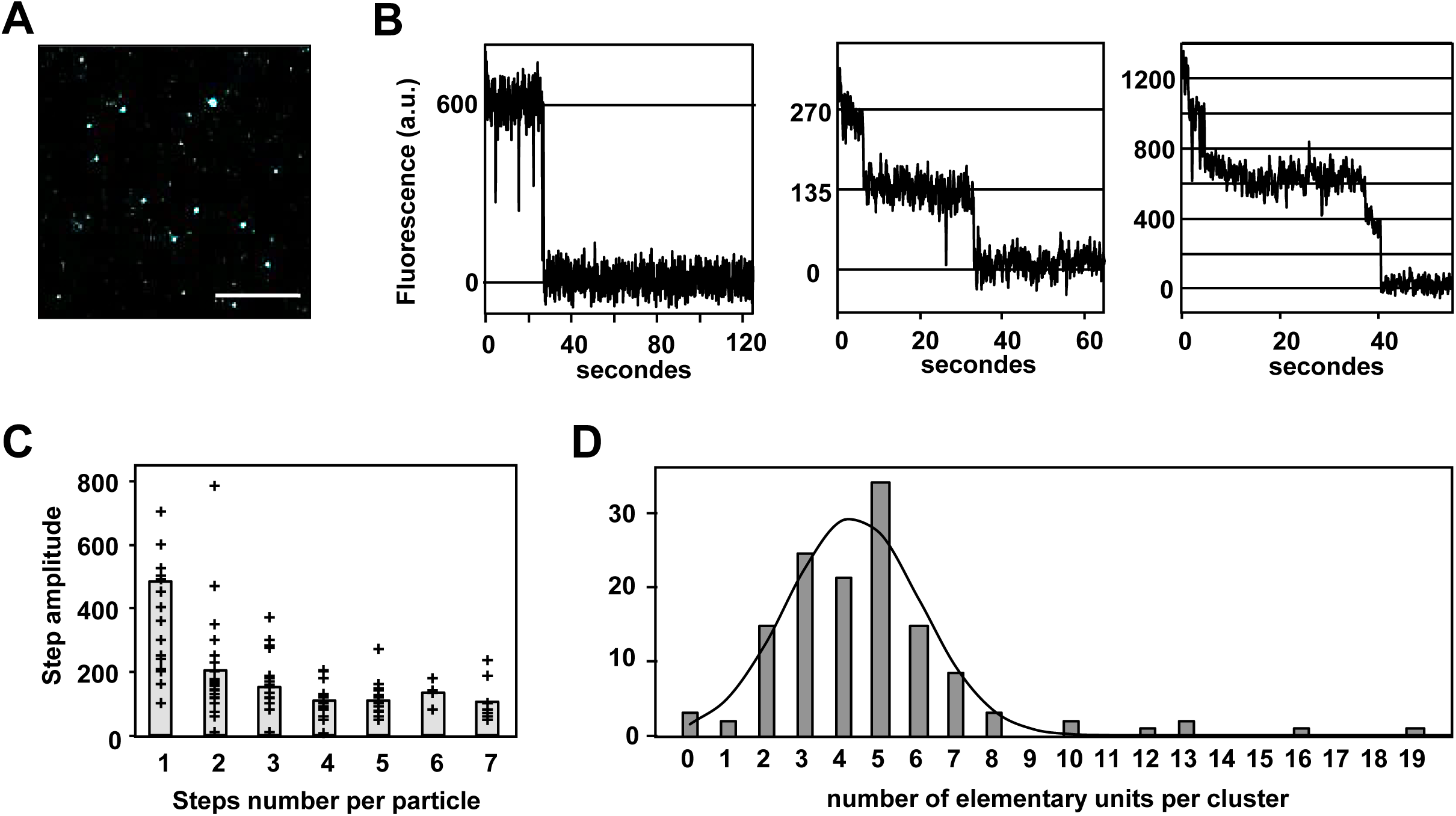
Single–particle fluorescence analysis of CX3CL1. A. Fluorescence of the membrane preparation of L_CX3CL1_ diluted in DOPC analyzed by TIRF. Bar 10µm B. Examples of the fluorescence of two particles tracked during two minutes C. Distribution of the number of elementary fluorescence units per particle calculated after analysis of the fluorescence kinetic of 126 different particles. The Gaussian curve fits the data with an amplitude of 27.5, a mean of 4.3 and a standard deviation of 1.8.

### Oligomerization of the peptide corresponding to the TM domain of CX3CL1

As evidenced by BRET (Hermand et al, 2008) and by FRAP (Ostuni et al, 2014), the monomers association of the membrane CX3CL1 molecule is mainly controlled by their transmembrane domain. We intended to directly investigate the possible aggregation of the TM peptide alone to avoid possible interferences of cytosolic and extracellular CX3CL1 domains. To this end, the peptide corresponding to the 20 residues of the TM domain was synthesized, with 2 lysine added at each of its end to ensure solubility. The resulting peptide is called TM24. As a control, we constructed a peptide called SCR24 (“scrambled”) with the same residues randomly rearranged, except the two lysines at each end. We first checked that these peptides fold correctly as alpha helices in appropriate buffer: when solubilized in trifluorethanol (TFE) or detergent, both TM24 and SCR24 possess more than 55% of helical structure, and more than 90% when solubilized in dodecylphosphocholine (DPC) for TM24 (Fig EV2). The conformation of TM24 in DPC micelles was further examined at the residue level by NMR spectroscopy. Analysis of NMR data confirms the helical conformation of the peptide, starting at residue Val3 and stabilized by numerous van der Waals contacts between Val, Leu and Phe side chains (Fig EV3).

We investigated the potential aggregation of the TM24 and SCR24 peptides by FRAP after peptides incorporation in Giant Unilamelar Vesicles (GUVs) of egg phosphatidyl choline (EggPC). To this end, the peptides were conjugated with fluorescein (FITC). Using bleaching circle spot of various diameters (2 to 10 µm), we found different recovery times (Fig 3A). The calculated diffusion rate is 0.09 ± 0.02 µm^2^sec^−1^ for TM24 and 1.14 ± 0.40 µm^2^sec^−1^ for SCR24, showing that TM24 diffuses considerably slower than SCR24 (Fig 3B), and hence is probably more aggregated than SCR24. This experiment was performed in GUV environment, i.e. a strict two-dimensional medium. We also used a tridimensional medium consisting of a lipid cubic “sponge” phase made with C_12_E_5_ lipids in the presence of n-octyl-β-D-glucopyranoside (Adrien et al, 2016; Pincet et al, 2016). In this case, the bleaching pattern was made with interference fringe (Davoust et al, 1982; Gambin et al, 2006; Reffay et al, 2009). A very similar result was found (Figs 3C and 3D). Since the very slow lateral diffusion of TM24 peptide – as compared to its SCR24 analogue - was observed in two different types of pure lipids structure, it most probably corresponds to an oligomerization due to intrinsic features of the peptide.

**Fig 3.**
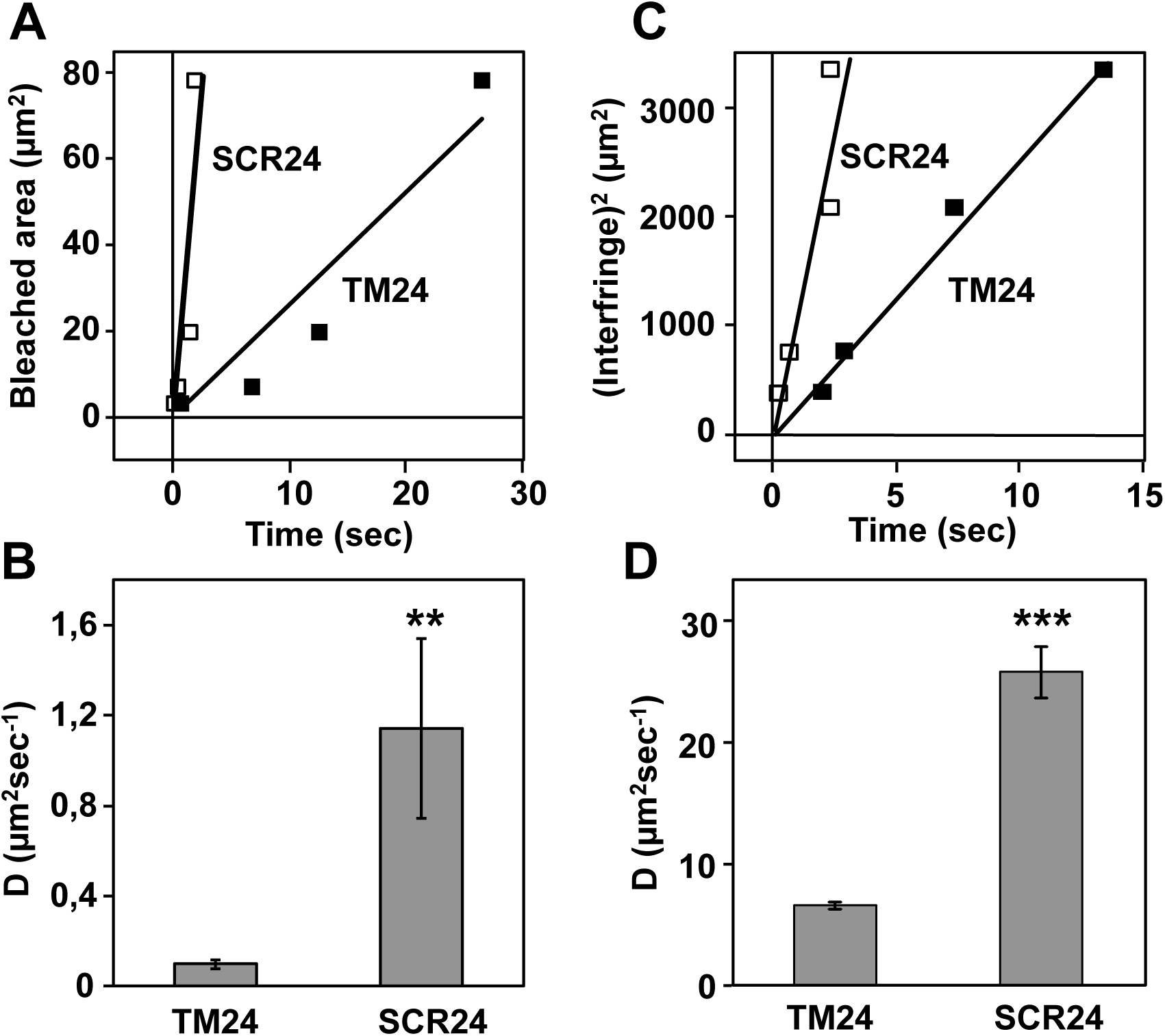
Diffusion rate of CX3CL1 transmembrane peptides analyzed by FRAP and FRAPP. A. The fluorescence kinetics of Giant Unilammelar Vesicles containing either TM24-FITC (filled squares) or SC24-FITC (empty squares) was analyzed after bleaching by circles on various diameter (1, 2, 5 and 10 µm). The recovery constant time was reported *versus* the bleached area. Each measurement was the mean of triplicates. The slopes of the linear fit are 2.63 and 29.93 for TM24 and SCR24 respectively. B. The diffusion rates were calculated based on the mean±SEM of the 12 measurements data used to give the Fig A. C. The fluorescence kinetics of lipidic cubic “sponge” phase containing either TM24-FITC (filled squares) or SC24-FITC (empty squares) was analyzed after bleaching by interference pattern with various interfringe (20, 27, 46 and 58 µm). The recovery constant time was reported *versus* the interfringe distance. Each measurement is the mean of triplicates. The slopes of the linear fit are 252 ± 10 and 1093 ± 144 for TM24 and SCR24 respectively. D. The diffusion rate were calculated based on the mean±SEM of the 12 measurements data used to give the Fig C. Note that the diffusion rate appeared dramatically higher in sponge phase since the fluorescent molecules could move in tridimensional milieu.

To directly catch its oligomerization, the peptides are subjected to crosslinking using bis(sulfosuccinimidyl)suberate (BS3) after dissolution in DPC, and then analyzed by SDS-PAGE (Fig 4A), as previously shown for the transmembrane domain of Carnitine palmitoyltransferase 1 (Jenei et al, 2009). In absence of crosslinker, TM24 and SCR24 bands appear between 3 and 6 kDa, i.e. as mixture between monomer and dimer (Fig 4A, left). After crosslinking, SCR24 displays as a large smearing spot from 30 kDa to 200 kDa probably corresponding to nonspecific aggregates (Fig 4A, right), which are also visible in the presence of SDS (Fig 4A, left). In contrast, TM24 shows discrete bands stepping from 3 to 26 kDa (Fig 4A), i.e. with 1 to 10 monomers, as revealed by densitometry (Fig 4B). This confirms that the TM24 peptide alone can correctly oligomerize.

**Fig 4.**
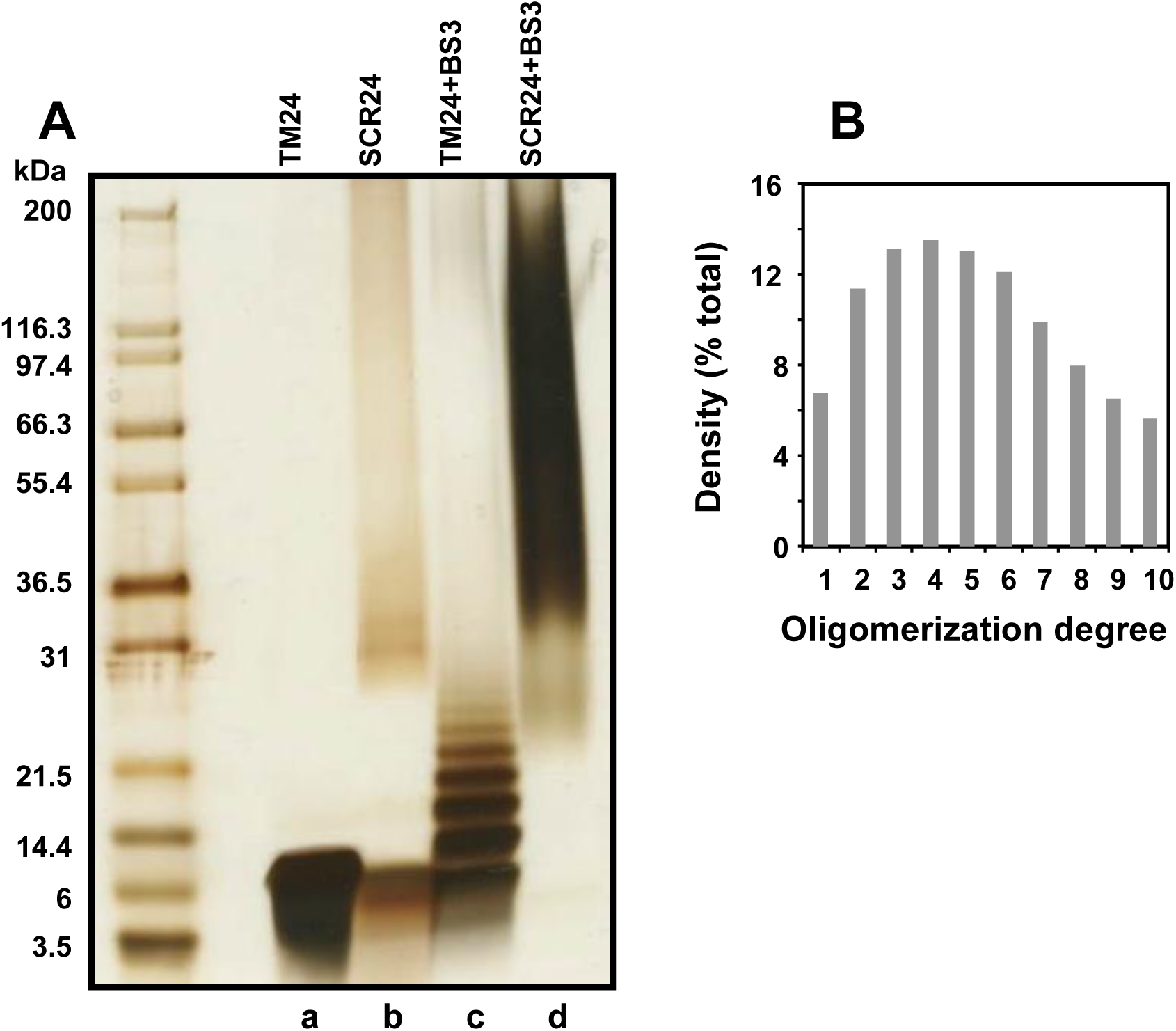
TM24 peptide polymerization analyzed by cross-linking and SDS-PAGE. A. SDS-PAGE of the TM24 and SCR24 peptides dissolved in DPC with a ratio [DPC]/[peptide] = 10 and cross-linked or not with 1.9 mM SB3 crosslinker. The gel is then silver stained. The left lane contains markers of various molecular weight. B. Densitogram of the “TM+SB3” lane in A.

The dynamical auto-association of TM24 has been assessed by coarse-grained molecular dynamics simulations. Three simulations in lipid boxes made of DOPC were built with one NMR structure of TM24. 9 or 16 monomers were included in DOPC boxes at different TM24/DOPC molar ratios (1/50 or 1/100) (Fig EV4A). The kinetics of TM24 peptide assembly followed over 10 µsec simulations revealed the formation of various types of oligomers ranging from dimers to nonamers (Fig 5A), depending on the initial conditions (Fig EV4B and EV4C). Once formed, the monomer to monomer interaction was stable over time leading to a compact interface, with an average distance between helix-helix center of mass lower than 1 nm (Fig EV4D). The dimer association did not occur at random, but a favored interface was observed where the TM are in almost parallel conformation, i.e. with azimuthal angle of 20-30° (Figs 5B and EV4C). Further oligomer association from this initial dimer showed an unexpected linear alignment of the TM interacting side by side (Fig 5A).

**Fig 5.**
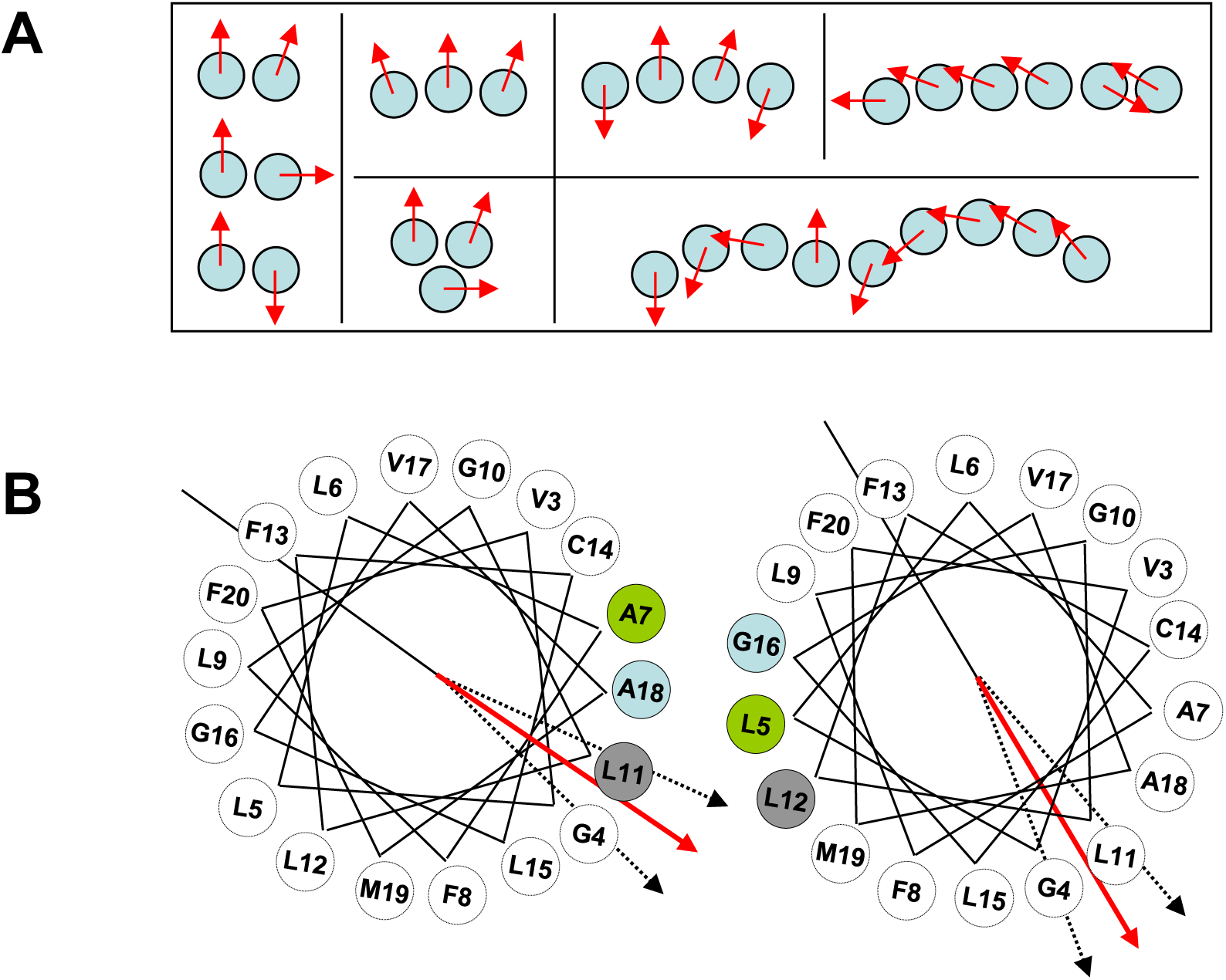
Oligomer formation observed by coarse-grained molecular dynamics simulations of TM24 in DOPC lipids. A. Topology of the main oligomers observed in the three simulations. Red arrow within circle indicates an arbitrary axis in the middle of the TM24 helix enabling to distinguish between parallel, orthogonal and symmetrical dimers. B. Helical wheels represent the most frequent dimer (parallel) interface involving Ala7, Leu11, Ala18 and leu5, Leu12, Gly16 from each monomers, respectively. Dotted arrows illustrate the angular fluctuation observed both for one dimer along the simulations and between the different dimers within this parallel arrangement.

Finally, the oligomerization degree of the TM24 oligomer was investigated in cellular context (Fig 6). The diffusion rate of the peptide expressed as a EYFP fusion in COS cells was compared with the ones of different transmembrane proteins that are known to oligomerize through their TM domains with a given number of monomers: the CD2 that is known to be monomer (Bodian et al, 1994; Jones et al, 1992), the transmembrane domain of the dimerized GlycophorinA (TM-GyA) (Lemmon et al, 1992), the pentamer phospholamban (Arkin et al, 1994; Colyer, 1993; Jones et al, 1985), the 7-TM CX3CR1 and the 12-mer RhBG (Callebaut et al, 2006). The latter was previously deglycosylated to ensure that the extracellular domain was not limiting the diffusion rate of the protein (Ostuni et al, 2014; Scullion et al, 1987; Wier & Edidin, 1988). We find that TM24 diffusion rate is close to the one of the CX3CR1, meaning that TM24 behaves as a heptamer in this cellular context. Taken together, our data performed with three different types of experiments (FRAP in pure lipid environments, SDS-PAGE after chemical cross-linking, FRAP in cellular context) all indicated the oligomerization of the TM24 peptide, with an oligomerization degree between 4 and 7.

**Fig 6.**
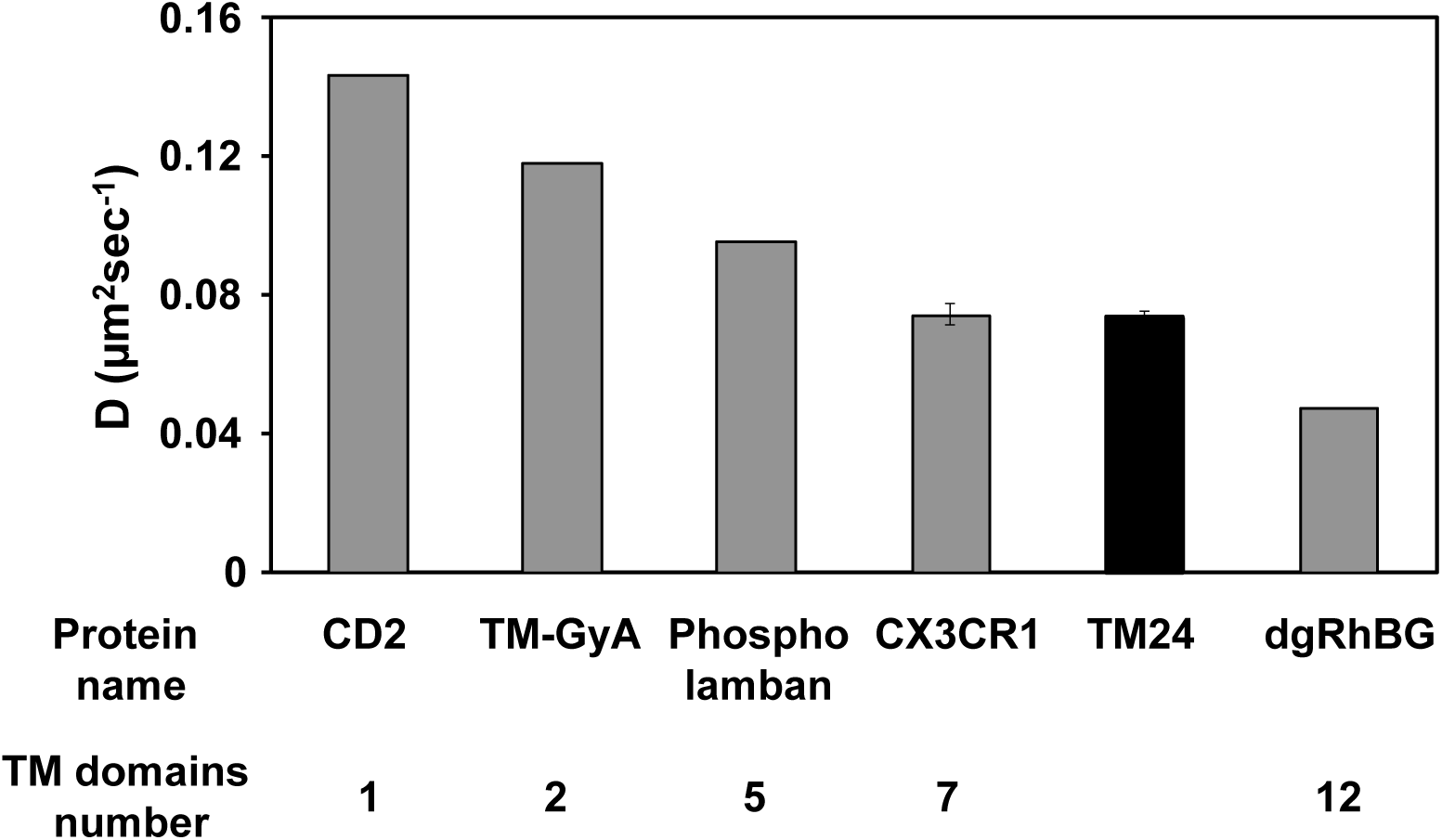
Diffusion rate in cellular membrane of TM24 peptide and various proteins with a known number of TM domains. The lateral diffusion rate of the TM24-FITC peptide and of other proteins with known TM number were assayed by FRAP after transitory expression in COS-7 cell line. Each point is the mean of duplicates, except for CX3CR1 and TM24 (mean of pentaplicates±SEM).

### Use of TM-CX3CL1 peptide to alter the CX3CL1 function

Our results strongly suggest that the CX3CL1 oligomerization is only controlled by the CX3CL1 TM domain. Therefore, the addition of peptide corresponding to the TM domain may compete with native CX3CL1 in oligomers, reducing the local concentration in complete chemokine and thereby altering the adhesive function of CX3CR1/CX3CL1 couple.

To directly test this hypothesis, we added the TM24 peptide to cells expressing the CX3CL1-EYFP fusion protein and analyzed its diffusion behavior by FRAP in COS-7 cells (Fig 7). Yet, we previously showed that CX3CL1 diffusion is highly controlled by the presence of the mucin stalk. If, as hypothesized, the presence of TM24 peptides generates CX3CL1 assemblies with a smaller amount of CX3CL1 monomers, the viscous drag due to extracellular mucin stalks will decrease, corresponding to an enhanced diffusion rate. This was exactly what we found: in the presence of 3 µM or 10 µM of TM24, the diffusion rate of CX3CL1 was considerably enhanced (Fig 7, filled bars). As a control, the addition of the SCR24 peptide had no effect (Fig 7, empty bars).

**Fig 7.**
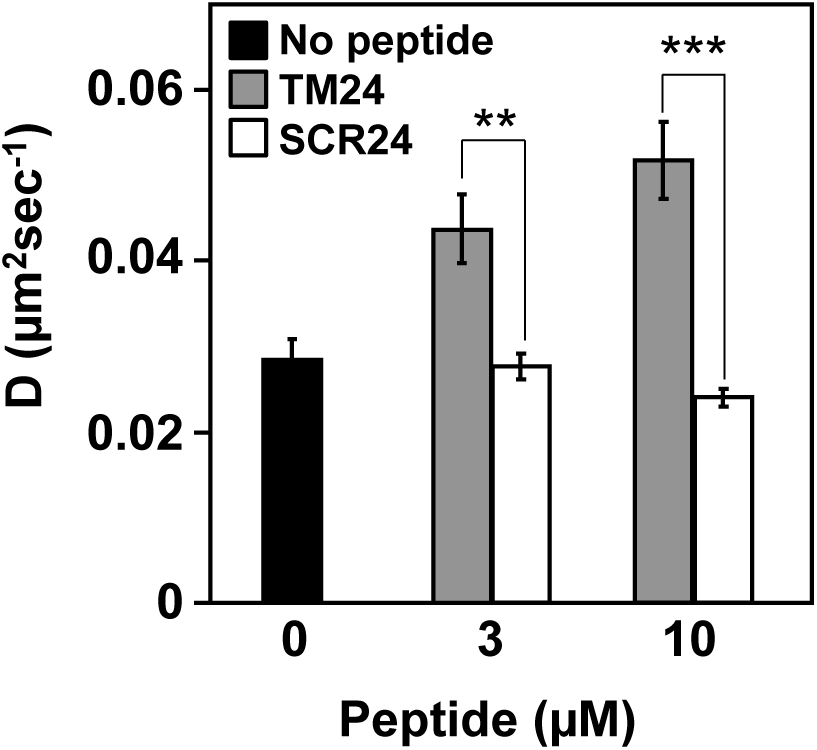
Diffusion rate in cellular membrane of the CX3CL1 protein in the presence of TM24 and SCR24 peptides. The lateral diffusion rate of the CX3CL1-EYFP protein was assayed after transient expression in COS-7 cell line after 15 min preincubation in the presence or not of 3 or 10 µM of TM24 or SCR24 peptides. Each point is the mean of triplicates±SD.

Having shown that TM24 peptides altered the CX3CL1 lateral diffusion by possibly dispersing the optimal CX3CL1 oligomer, we measured the impact of TM24 on the adherence function. To this end, we quantified in real time the cell to cell adherence using the LigandTracer™ technique (Hillerdal et al, 2016). As controls, the black solid trace of Fig 8A represented the adherence of CHO expressing CX3CR1 (here called CHO_CX3CR1_) toward L_929_ cells expressing CX3CL1 (called here L_CX3CL1_), while the black dashed trace represented the adherence of the CHO_CX3CR1_ to non-transfected L_929_ cells. The adherence to L_CX3CL1_ was 3.5 fold higher than the non-specific adherence to L_929_. In the presence of 5 µM TM24, the CHO_CX3CR1_ slightly more adhered to L_929_ (Fig 8A, red dashed trace). By contrast, the CHO_CX3CR1_ adherence to L_CX3CL1_ was dramatically reduced (more than 2.2 fold less) (Fig 8A, red solid trace) to the level of the non-specific adherence to L_929_ (compare both red traces in Fig 8A). In the presence of 5 µM SCR24, the CHO_CX3CR1_ adherence to L_CX3CL1_ was unchanged (Fig 8A, blue solid trace) while the adherence to L_929_ was similar to that in the presence of TM24 (Fig 8A, blue dashed trace). Then, we calculated the specific adherence, i.e. the adherence to L_CX3CL1_ subtracted by the adherence to L_929_ (Fig 8B). The inhibition of the CX3CL1-specific adherence of the CHO_CX3CR1_ cells by TM24 was nearly complete (compare the red trace to the control black trace). This inhibition was dependent on the peptide sequence since the scrambled one had almost no effect (blue trace, Fig 8B). This result was further borne out by showing the result of triplicate experiments after 60 min of adherence (Fig 8B right): 5 µM TM24 reduced the CX3CL1-specific adherence to zero, while 5 µM SCR24 restrained it only by 20%.

**Fig 8.**
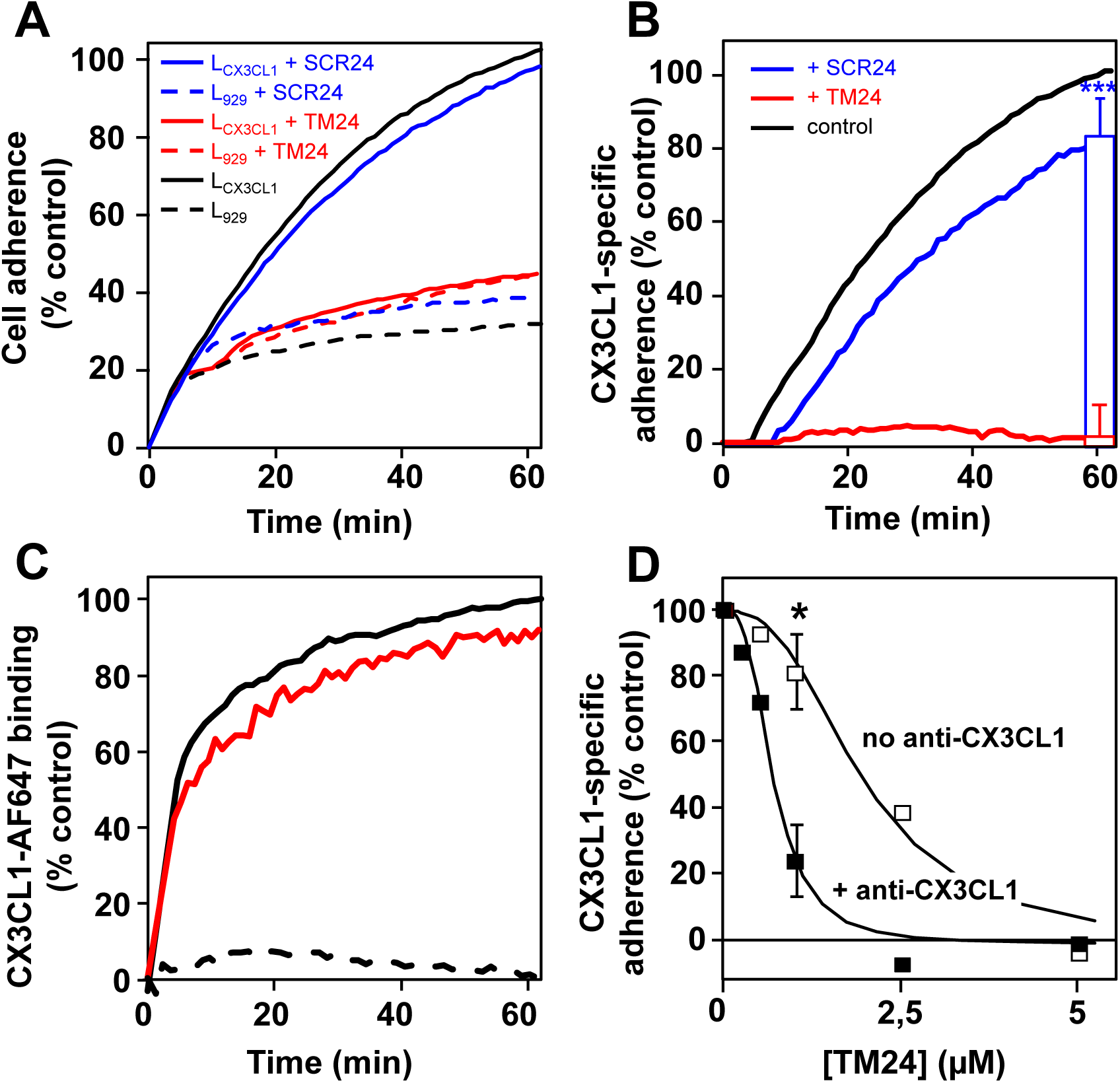
Specific inhibition of the CX3CL1-dependent cell adherence by the peptide TM24. A. Real time adherence of CHO_CX3CR1_ cells to L_929_ (dashed traces) or L_CX3CL1_ cells (solid traces) as assayed by the LigandTracer™ technique, in the presence of 5µM TM24 (red traces), 5µM SCR24 (blue traces) or none (black traces). The data were normalized using the control trace with L_CX3CL1_ cells in the absence of peptide (100% after 60 minutes). The curves are the mean of three independent experiments. B. Specific adherence of CHO_CX3CR1_ cells to L_CX3CL1_ cells using data of Fig 7A. The data obtained with L_CX3CL1_ cells were subtracted from data obtained with L_929_ and normalized at 100% at the 60 minutes time. The bars in the right of the Fig shows the specific adherence after 60 minutes (mean of triplicates±SD) in the presence of 5µM TM24 (red) and 5µM SCR24 (blue). C. Real time binding of 100nM of fluorescent CX3CL1 to coated CHO_CX3CR1_ cells using the LigandTracer™ technique, in the presence of 5µM TM24 (red trace), 1µM unstained CX3CL1 (dashed trace) or none (black trace). The data were normalized using the control trace without peptide (100% after 60 minutes). D. Specific adherence of CHO_CX3CR1_ cells to L_CX3CL1_ cells after 60 minutes in the presence of various TM24 peptide concentrations and in the presence (filled squares) or in absence (empty squares) of 1µg/ml of anti-CX3CL1 antibody. The data were calculated and normalized as in Fig 7B. Experiments are performed in duplicate except for the 1µM TM24 concentration done in triplicates (mean±SD).

To check that this TM24 specific effect is not due to some interference with CX3CL1-CX3CR1 binding interface, we conducted binding experiments using the fluorescent CX3CL1-AF647 with the same LigandTracer™ technique (Fig 8C). In the presence of the TM24 peptide, the binding of the chemokine to the CHO_CX3CR1_ was only marginally affected (Fig 8C, red trace), while it was completely suppressed in the presence of an excess of unlabeled CX3CL1 (Fig 8C, dashed trace). Next, we assayed CHO_CX3CR1_ calcium response to soluble CX3CL1 (Fig EV5): the presence of the TM24 peptide had no effect, confirming that the presence of the cellular transmembrane CX3CL1 molecule is needed to reveal the TM24 action. Finally, we performed the cell-to-cell adherence assay in the presence of anti-CX3CL1 antibodies (clone AF365), which are known to bind to CX3CL1 surface directly binding to CX3CR1. Indeed, this anti-CX3CL1 antibody clearly inhibited the CHO_CX3CR1_ adherence to L_CX3CL1_ (Fig EV6). In the presence of a concentration of this anti-CX3CL1 that only marginally alters the CHO_CX3CR1_ adherence (1 µg/ml), we found that the dose-response curve of the TM24 peptide significantly shifted to the left (Fig 8D) meaning that less peptide was required to obtain the same inhibition. The presence of 1 µM TM24 was only slightly inhibitory without antibodies, while it almost wholly inhibited the specific adherence of CHO_CX3CR1_ cells in the presence of 1 µg/ml of antibodies (Fig 8D). Taken together our data showed that TM24 peptide effectively weakens the functional CX3CL1-CX3CR1 interaction by acting on the transmembrane domain of the CX3CL1 molecule.

## Discussion

The cell-to-cell adhesion mediated by CX3CL1-CX3CR1 interaction very probably implies numerous molecules at each cellular adhesive interface. Indeed, the membrane form of CX3CL1 self-associates in many cellular contexts, with an important role of the TM domain as shown by our previous works (Hermand et al, 2008; Ostuni et al, 2014). However, the question remained open to know if this association was an intrinsic property of the chemokine molecule or is due to an interaction with another molecular partner keeping the CX3CL1 monomers together. We therefore conducted here experiments with the TM24 peptide alone. As expected, this peptide is structured as an alpha-helix in lipid environment (Figs EV2 and EV3). Moreover, we show that the lateral diffusion rate of the TM24 peptide is considerably higher than its SCR24 analog in pure lipidic context, as shown by FRAP experiments using two completely different lipid structures (Fig 3). This strongly indicates that TM24 self-association is a multimerization due to hydrophobic forces that are specific of the sequence of the TM domain of CX3CL1. Electrophoresis data (Fig 4) confirm that the peptide assembly is properly organized since cross-linking reveals discrete bands with 1 to 10 monomers. We cannot exclude that the peptide oligomer contains some lipid inside, and that some lipid moieties are mandatory for oligomerization; however, all the lipids we used here (DOPC, Egg-PC, Figs 2 and 3A) can endow this organizing function. It also persists in L3 sponge phase (Fig 3C) and in DPC micelles (Fig 4).

We therefore performed experiments with the whole CX3CL1 molecule. According to our gel electrophoresis data in the presence of mild detergents (Figs 1B and EV1), CX3CL1 molecule is present at least as a trimer; a similar result was obtained after solubilization using stabilizing Calixarene-based detergent and immune-purification (Figs 1C). This shows clearly that CX3CL1 does not exist as monomer but rather as high molecular weight populations. Moreover, we cannot exclude that some of the oligomers were dissociated due to the detergent and the pH shock of the immuno-purification. Furthermore, our single particle fluorescence analysis indicates an oligomerization degree between 3 and 7 (Fig 1). This experiment requires a high dilution in DOPC, meaning that the observed oligomerization is due to intrinsic properties of the CX3CL1 molecule. Here again, it is not impossible that the amount of monomers per particle could be underestimated since the EYFP fluorescence could weaken before performing the experiment.

So our data converge to bear out that the CX3CL1 molecule does not behave as a dispersed monomer, but as a polymer and at least as a trimer. We moreover cannot exclude that CX3CL1 polymers include diverse populations with different oligomerization degree. Based on all the present data (Figs 1-6), the CX3CL1 appears as oligomeric molecule containing between three and seven molecules. Morover, the *in silico* experiments based on NMR structure of TM24 confirm our *in vitro* and *in cellulo* data demonstrating that oligomerization is an inherent property of CX3CL1-TM domain, even if the coarse-grained Martini model used in the molecular dynamics calculations tends to generate excessive aggregation of transmembrane helices (Javanainen et al, 2017).

Our FRAP experiment is consistent with such conclusion since the diffusion rate of the TM24 expressed in cellular context is as slow as the 7-TM CX3CR1 (Fig 6). Molecular modelling suggests that the CX3CL1 oligomer is not organized as a bundle but rather as a linear assembly (Fig 5A), favored by an interface involving Leu, Ala and Gly residues (Fig 5B). 3 to 5 linearly arranged TM would give the same Stokes radius as 7 closed packed TM (see Materials and Methods). In any case, the CX3CL1 oligomerization allows to better capture the CX3CR1 molecules of the counter-adhesive cell, especially if the CX3CR1 is linearly organized to accommodate the potential dimer of CX3CR1 (Darbandi-Tehrani et al, 2010).

There are several adhesive molecules whose adhering function involves its homodimerization through their transmembrane domain as integrin, ICAM-1 (Hynes, 2002; Miller et al, 1995). Others are oligomeric like cadherin (Strale et al, 2015) or connexin (Herve et al, 2004) but in this case, the oligomerization is mainly driven by the extracellular domain and appears in specific membrane context (adherens or gap junctions) (Hong et al, 2010; Shapiro & Weis, 2009; Thompson et al, 2019). So, to our knowledge, CX3CL1 is the first example of an oligomeric adherence molecule with at least three monomers in any cellular and lipidic context.

The next question we address is to know if this TM-driven CX3CL1 oligomerization could be inhibited by adding TM24 peptides. We first performed FRAP experiments using CX3CL1-EYFP expressing cell line. The addition of micromolar concentration of TM peptide indeed lead to significantly increased lateral diffusion, meaning a lighter CX3CL1 probably due to decrease of its oligomerization degree (Fig 7). This prompted us to check if the CX3CL1 adherence function is dependent of its oligomerization. To this end, we used a new technique able to follow in real time the cellular adherence in a confined volume (Fig 8). We found that in the presence of 5 µM TM24 peptide, the CX3CL1-CX3CR1-dependent cell-to-cell adherence is wholly inhibited, while the presence of 5 µM SCR24 has no effect (Figs 8A-8B). To confirm that the TM24 peptide does not directly act on the CX3CL1-CX3CR1 binding, we show that the binding of soluble and fluorescent CX3CL1 to immobilized CX3CR1-expressing cells was unaffected by the peptide (Fig 8C). We also show that the peptide has no effect on the cell calcium response triggered by the soluble form of CX3CL1 (Fig EV5). This means that the TM24 peptide does not affect the CX3CR1 binding interface to CX3CL1, but rather acts by decreasing the number of functional CX3CL1 monomers in the CX3CL1 bundles able to efficiently interact with CX3CR1. This was further shown by using an anti-CX3CL1 antibody, known to interfere with the CX3CL1-CX3CR1 binding (Fig 8D). This antibody acts also by decreasing the number of CX3CL1 molecules able to interact with CX3CR1. As expected, in the presence of such antibody, the TM24 inhibiting effect was potentiated (Fig 8D). So the TM24 peptide could allosterically or heterotopically control the availability of CX3CL1 oligomers to induce potent cellular adherence, in a similar way to the integrin activating peptide corresponding to the integrin TM domain (Yin et al, 2006). It would be interesting to test shorter peptides targeting specific spots of CX3CL1 transmembrane domain, in order to get inhibitors with better affinity.

In conclusion, our work demonstrates that CX3CL1 is an oligomeric molecule with at least three monomers and that its oligomerization is required for its adhesive function. We also show that this functional oligomerization could be controlled by a specific peptide corresponding to the CX3CL1 TM domain that does not affect the function of the CX3CL1 soluble form. This could be useful to control the CX3CL1 function specifically linked to its adherence feature, as the monocytes margination in vasculature (Hamon et al, 2017) and the monocyte retention in bone marrow (Jacquelin et al, 2013). This tool could be of peculiar worth to reduce the atherosclerosis by reducing the monocyte/macrophage adherence to inflamed endothelia. It could also reduce the *Falciparum* gametocyte maturation by inhibit their nesting in bone marrow.

## Materials and Methods

### Chemicals, proteins and cell culture

Human CX3CL1 (Chemokine Domain) and polyclonal goat anti-CX3CL1 antibody (clone AF365) were purchased from Biotechne (Lille, France). Peptides (KKVGLLAFLGLLFCLGVAMFTYKK called TM24, KKTLVACLVFGMLGYLAGFL-FLKK called SCR24, TM24-FITC, SCR24-FITC) were synthetized either by the peptide synthesis facility of the Institut de Biologie Paris-Seine (FR3631, Sorbonne Université, CNRS) or by ProteoGenix (Schiltigheim, France). A human embryonic kidney cell line (HEK293), the Chinese Hamster Ovary cell line (CHO), the COS-7 cell line and the mouse connective tissue L_929_ cell line were grown in Dulbecco′s Modified Eagle′s Medium (DMEM) supplemented with 10% fetal calf serum (FCS), 1% sodium pyruvate, and antibiotics. Stable transfections with the pEYFP *cx3cl1* construct (Ostuni et al, 2014) were performed using JetPei (PolyPlus Transfection, Illkirch, France) according to the manufacturer’s instructions. Stably transfected cells were selected with 1 mg/ml geneticin (G418, ThermoFisher Scientific, les Ulis, France), and single clones were established by limited dilution.

### Cell membrane preparation for electrophoresis

L_929_ cells stably expressing CX3CL1-YFP (hitherto denoted L_CX3CL1_) were harvested from culture flasks through treatment with Cell Dissociation Buffer (Life Technologies, Thermo Fisher Scientific), washed in PBS, and centrifuged. The pellet was suspended in Lysis Buffer (Tris 10 mM pH 8) for 60 min at 4 °C. Cell lysis was performed on ice using a Bead Beater homogenizer with 0.1 mm diameter glass beads. Membrane fractionation was then carried out at 4 °C by sequential centrifugations. Three centrifugations were performed: 500xg for 5 min, 15000xg for 30 min, and 100000xg for 45 min. Membrane enriched pellets corresponding to plasma membranes (100000xg) were resuspended in PBS, 200 mM NaCl, 1X protease inhibitor cocktail and glycerol 10%, quantified using the BCA method (Pierce, Thermo Fisher Scientific, Courtaboeuf, France), flash-frozen and stored at −80 °C until use.

### SDS-PAGE

CX3CL1 samples (10µg) were denatured with 5x Laemmli buffer and incubated for 20 min at RT prior to analysis without heating to avoid aggregates formation. Proteins were separated by SDS-PAGE on a 4-15% acrylamide gel (4–15% Mini-PROTEAN® TGX Stain-Free™ Gel, *Bio-Rad*) and subsequently immobilized by electro-transfer to PVDF membrane.

### Native PAGE Native PAGE of solubilized proteins using digitonin and dodecylmaltoside (DDM)

CX3CL1 samples were suspended in 75 µl of Native PAGE sample buffer (Thermo Fisher Scientific) in the presence of either 1% digitonin (Sigma) or DDM (Sigma) supplemented with Complete EDTA-free protease inhibitor (Roche) for 30 min at 4 °C under shaking. For proteins separation, 40 μg were loaded in NativePAGE Novex Bis Tris Gels (3-12 %) and transferred on a PVDF membrane according to the manufacturer’s instructions (ThermoFisher Scientific). Gels were electrotransferred to Hybond-P nitrocellulose membrane (Amersham Biosciences), and the blots probed with polyclonal goat antibodies anti-CX3CL1. For detection, we used horseradish peroxidase-conjugated goat anti-mouse IgG (Bio-Rad) and an enhanced chemiluminescence detection system (Amersham Biosciences).

### Clear Native-PAGE of calixarene based solubilization and immuno-purification

Proteins from plasma membrane fractions were incubated for 2 h at 4 °C at a final concentration of 2 mg/ml in 50 mM phosphate buffer pH 8.0, 200 mM NaCl, 1X protease inhibitor cocktail, 20 glycerol and with 5 Critical Micellar Concentration of CALX173ACE (CALIXAR). CALX173ACE solubilized CX3CL1 was loaded into magnetic beads previously crosslinked to polyclonal anti-CX3CL1 antibody using an IP kit (Pierce, Thermo Fisher Scientific). Retained CX3CL1 was eluted by pH shock (basic) followed by a neutralization step. Non-denaturated proteins were separated by native-PAGE on a 4-15% acrylamide gel (4–15% Mini-PROTEAN^®^ TGX Stain-Free™ Gel, *Bio-Rad*) using 25 mM imidazole pH 8.0 as anode buffer and 50 mM Tricine, 7.5 mM imidazole, 0.05% deoxycholate and 0.01% DDM as cathode buffer). Clear Native PAGE gels ran for 90 min at 200 V and 4 °C. Proteins were then immobilized by electro-transfer to PVDF membrane. The immunodetection of CX3CL1 was performed by using the SNAP i.d. system (*Millipore*) with primary antibody (R&D system, MAB3651) against CX3CL1 (1/500 dilution) and revealed using a mouse HRP secondary antibody (Santa Cruz, 3/10,000 dilution).

### Single-molecule photobleaching TIRF

L_CX3CL1_ clone cells were harvested from culture flasks through treatment with Cell Dissociation Buffer (Life Technologies, Thermo Fisher Scientific), washed in PBS, and resuspended in Lysis Buffer (Tris 10 mM pH 8, 2 mM EDTA) supplemented with Complete protease inhibitor (Roche, Merck, Sigma-Aldrich, l’Isle d’Abeau, France). After three cycles of freezing – thawing, the lysate was centrifuged (500xg, 5 min, 4 °C). The supernatant was centrifuged at 4 °C at 16000xg during 45 min. The pellet containing the membrane was then resuspended in the lysis buffer. 500µl of this pellet suspension was added to a tube in which 2.5 mg of 1,2-dioleoyl-sn-glycero-3-phosphocholine (DOPC, Avanti Polar lipids, Merck, Sigma-Aldrich, l’Isle d’Abeau, France) in chloroform had previously been evaporated. The tube was vortexed for 45 min. 10 µl of this mixture was deposited on a coverslip. Membranes from homogenates and DOPC liposomes spontaneously spread on the coverslip. The concentration of homogenates was low enough to observe individual fluorescent clusters separated by large non-fluorescent DOPC regions. The particle fluorescence was analyzed by an inverted microscope (Eclipse Ti, Nikon Instruments, Amsterdam, Netherlands) using a 60x oil objective. The excitation light is provided by a 488 nm laser beam with an incidence larger than the critical angle to ensure total reflection. For each monitored region of the coverslip, a 2 min movie was recoded. Bleaching of the clusters was subsequently analyzed using ImageJ.

### FRAP in pure lipid environment with giant unilamellar vesicles (GUVs)

GUVs containing peptides were formed using the osmotic shock protocol (Motta et al., DOI: 10.1021/acs.langmuir.5b01173, Langmuir 2015, 31, 7091−7099). In brief, 300 µg of EggPC (L-α-phosphatidylcholine (Egg, Chicken), Avanti Polar Lipids) were dried in a glass tube. In parallel, TM24-FITC and SCR24-FITC were solubilized at 1.2 µM in PBS. 330 µl of each solution was added in different EggPC containing tubes that were subsequently vortexed for an hour to form liposome-like structures containing 1000 lipids per peptide. On Mattek dishes (Mattek Corporation), 2 µl of each solution was dried and rehydrated with 2 µl of pure water (3 cycles) to form GUVs. After the last cycle, the Mattek dishes were filled with 500 µl PBS. FRAP experiments were performed using a Leica 63x dry objective. Two identical regions of interest (ROI) were monitored: one was photobleached during three scans with the 488 nm laser beam at full power, and the other was used to monitor the concomitant effects of intrinsic photobleaching. The pre- and post-bleach images were monitored at low laser intensity (10 to 15% of full power). The fluorescence in the ROIs was quantified using the LASAF Leica software. The analysis of the curve of the resulting fluorescence recovery as a function of time yielded the recovery times that were used to obtain the diffusion coefficients of the diffusing species. The diffusion coefficient is equal to D = r^2^/4τ where “r” is the radius of the circular beam and “τ” is the time constant obtained from the fit of the curve (Soumpasis, 1983). In our experiments, each cell was bleached over three different circular regions with diameters of 3, 4, or 5 µm. The characteristic recovery times (τ) and fluorescence recovery plateau (Fp) were calculated by fitting the fluorescence recovery curves as previously described (Braeckmans et al, 2003; Soumpasis, 1983).

### Fluorescence recovery after pattern photobleaching

Fluorescence recovery after pattern photobleaching (FRAPP) experiments were performed as previously described (Adrien et al, 2016; Rayan et al, 2010). Briefly, TM24-FITC or SCR24-FITC peptides were added into a L3 sponge phase prepared by mixing the non-ionic surfactant C_12_E_5_ (pentaethylene glycol monododecyl ether), the co-surfactant β-octylglucopyranoside and PBS. The samples were vortexed, sonicated, and equilibrated at room temperature for at least 10 min. The distance among the bilayers is tuned by varying the amount of these 2 compounds as previously described (Reffay et al, 2009)

The L3 phase was injected into capillary tubes of a 200 μm thickness (VitroCom, Mountain Lakes, New Jersey), sealed with wax in order to prevent evaporation. Two interfering laser beams were focused on a dot of approximately 250 μm of diameter creating a fringe pattern. The interfringe distances were tuned from i = 19.3 to 44,2 μm. The recovery curves were fitted by an exponential decay. In all experiments, we observed a pure monoexponential recovery of fluorescence, indicating the diffusion of monodisperse objects. The fact that the diffusion was Brownian was verified by obtaining a linear variation of the characteristic recovery times τ plotted versus i^2^, and the D value was deduced from the slope of the linear regression following this equation: D = i^2^/4π^2^τ. The diffusion coefficients were measured with an accuracy of 10%. Experiments were performed at room temperature.

### FRAP in cellular environment

COS-7 cells were seeded into four wells borosilicate Labtek chambers (Costar-Corning, NY, USA) 48 h and transfected 24 h before the FRAP experiments. Confocal imaging was performed on live cells with a Leica SP5 microscope using a 488-nm laser beam and the filter sets supplied by the manufacturer. The cells were maintained at 37 °C on the microscope stage and FRAP experiments we performed in a similar manner as for the GUVs using three different circular bleached regions with diameters of 3, 4, or 5 µm.

### Peptide crosslinking and electrophoresis

Peptide crosslink in detergent was performed as previously described (Jenei et al, 2009). Briefly cross-linking reactions were carried out for 0.4 mg/ml solutions of the SCR24 or TM24 peptide dissolved in dodecylphosphocholine (DPC) micelles (DPC/peptide = 0.1). All samples were prepared in 20 mM sodium phosphate buffer and 150 mM NaCl (pH 8). Bis[sulfosuccinimidyl]-suberate (BS3, Pierce, 1.9 mM final) was used to cross-link the peptide according to the manufacturer’s protocol. The cross-linking reaction was stopped after 30 min by the addition of 1 M Tris-HCl (pH 8). Samples were separated by SDS-PAGE and revealed by staining with silver nitrate.

### Circular dichroism

Far UV circular dichroism spectra were recorded on a Jobin-Yvon CD6 spectropolarimeter operated at room temperature according to published protocols (Jamin & Lacapere, 2007). Briefly, a peptide sample (5 µM) in 10 mM sodium phosphate buffer solution without or with 30% TFE (TriFluoroEthanol) and 80 mM SDS or DPC was placed in a 0.2 mm path length quartz cuvette (Hellma, France). CD spectra were recorded in the 185 to 270 nm wavelength range with a 0.2 nm step resolution, 1 s signal averaging time and 1 nm bandwidth. Spectra were averaged over five scans, corrected for background and smoothed over 25 points. A consensus secondary structure content was estimated by spectral deconvolution using CONTINLL, CDSSTR and Selcon software and data sets of reference proteins SMP50 (37 soluble proteins and 13 membrane proteins) and SP37 (37 soluble proteins) (Sreerama & Woody, 2000), as well as CDFriend.

### NMR

TM24 was dissolved at 1 mM concentration in 550 μL of H_2_O/D_2_O (90:10 v/v) containing 80 mM DPC. NMR experiments were acquired on a Bruker Avance III 500 MHz spectrometer equipped with a TCI ^1^H/^13^C/^15^N cryoprobe. A double pulsed field gradient spin-echo was applied by using band-selective shaped pulses (90° G4 pulse of 4 ms, 180° Reburp pulse of 3 ms) centered on the amide/aromatic region so as to eliminate large ^1^H signals arising from non-deuterated detergent in the acquisition dimension. NMR spectra were processed with TopSpin 3.2 software (Bruker) and analysed with NMRFAM-SPARKY program (Lee et al, 2015). ^1^H resonances were assigned (Table EV1) using 2D ^1^H-^1^H TOCSY (DIPSI-2 mixing time of 68 ms) and 2D ^1^H-^1^H NOESY (150 ms mixing time) recorded at 40°C. The chemical shift deviations of ^1^Hα were calculated as the differences between observed chemical shifts and random coil values reported in water (Wishart et al, 1995). 188 inter-proton distance restraints (71 intraresidual, 85 sequential, 32 medium-range) were derived from NOESY cross-peak intensities and 43 dihedral angle restraints (22 ϕ, 21 ψ) were obtained from the analysis of ^1^Hα CSDs. Structures were calculated using Amber 14 program (Case et al, 2014) and ff14SB force field (Maier et al, 2015), as described (Byrne et al, 2019).

### Molecular dynamics

Coarse-grained (CG) molecular dynamics simulations were carried out using the MARTINI force field for lipids (Marrink et al, 2007) and proteins (Monticelli et al, 2008). Both electrostatic and Lennard-Jones interactions were calculated using a 1.2-nm cutoff with a switch function. The neighbor list was updated every 20 steps and the relative dielectric constant for the medium was set to 15. The temperature for each group (peptide, lipid, water) was kept at 303.15 K using the v-rescale temperature coupling algorithm (Bussi et al, 2007) with a time constant of 1 ps. The pressure was kept constant using the Parrinello-Rahman algorithm (Parrinello & Rahman, 1981) with a semiisotropic pressure coupling (*x* and *y* dimensions, the bilayer plane, coupled independently from the *z* dimension) and a time constant of 12 ps. The integration time step was 20 fs and structures were saved every 100 ps for analysis.

Three boxes containing 9, 9 and 16 TM24 at lipid ratio 1:100, 1:50 and 1:100 (respectively) were constructed (Box1, Box2 and Box3) using CHARMM-GUI (Jo et al, 2008; Qi et al, 2015). At the beginning, we placed one TM24 (using the best NMR structure) in a vertical position within a bilayer of dioleoylphosphatidylcholine (DOPC) (of 100 lipids for Box1 and Box3 and 50 lipids for Box2. The system was hydrated with slightly more than ten times more water beads compared to the number of lipids (corresponding to an effective water to lipid ratio of 1:40, one water bead corresponds to 4 water molecules). We also added Na^+^ / Cl^−^ ion beads to get an ion concentration of ~150 mM and keep the system neutral. This system was energy minimized and equilibrated to pack the lipids on the peptide (using position restraints on the latter). Then, the system was replicated 3 times (for Box1 and Box2) or 4 times (for Box3) on the *x* and *y* directions using the *genconf* program from the GROMACS package. After a minimization and an equilibration of 10 ns (with position restraints on the peptide), production simulations were run for 10 μs for each Box. All simulations were carried out using GROMACS 2018.5 (Abraham et al, 2015).

### Stokes radius of a linear oligomer

To our knowledge, there is no model for the 2D-hydrodynamic radius (Stokes radius) of linearly aligned transmembrane domains in a membrane. Qualitatively, a compact disc-like structure will have a smaller 2D-hydrodynamic radius than an elongated rectangle with the same area. To have a more quantitative insight, we propose to model the linear arrangement of n_r_ CX3CL1 as an elongated rectangle with 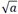 as a width and 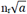 as a length; *a* is the area occupied by a transmembrane domain. The stokes radius can be estimated by the average projection of the rectangle in all directions: (n_r_ + 1)*a*/π.

In a cellular context, we found TM24 diffuses as fast as CX3CR1 that has seven transmembrane domains. Assuming the arrangement of CX3CR1 transmembrane domains has a disk-shape, the 2D-hydrodynamic radius is 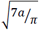. Hence, the number of CX3CL1 in a linear arrangement with the same 2D-hydrodynamic radius would be n_r_ ~ 4.

### Calcium flux assay

A cytosolic-free calcium assay was measured by fluorescent detection using CalbryteTM520-AM (AAT Bioquest, Interchim, Montluçon, France), according to the manufacturer’s instructions. In brief, CHO-CX3CR1 cells were plated overnight (4×10^4^ cells per well) in black 96-well microplate with clear bottom (Greiner, Dutscher, Brumath, France) at 37 °C, 5% CO_2_. Then the cells are loaded for 45 min at 37 °C with a solution containing 0.04% pluronic acid (Interchim), 10 µM Calbryte™520 AM and 1 mM probenecid (Interchim) in working buffer called HHBSS (Hank’s Balanced Salt Solution supplemented with 10 mM Hepes, 1 mM MgCl_2_ and 1 mM CaCl_2_, pH 7.2). After washing, the cells were treated with control buffer or TM24 peptide and then with soluble CX3CL1 (Chemokine Domain) at the appropriate concentrations. Signal was measured as a function of time on a fluorescent plate reader equipped with fluidic handling (FlexStation 3; Molecular Devices, Sunnyvale, CA, USA) and quantified by SoftMax Pro software (Molecular Devices). The signal of each well was normalized using the maximum obtained after addition of ionomycin (Interchim) and the minimum obtained after addition of EGTA.

### Cellular adherence assay using LigandTracer™

The day before the experiment, 5×10^5^ L_929_ or L_CX3CL1_ cells were seeded in 35 mm Petri dishes (Cellstar Greiner, Dutscher, Brumath, France). After washing, a volume of 900 µL of HHBSS was kept in each dish. Four of such dishes were inserted in an 87-89 mm cell dish, and the whole setup was placed on the tilted and rotating support of the LigandTracer instrument (Ridgewiew, Vänge, Sweden) equipped with the 488/535 detector. The fluorescence intensity in the elevated area - i.e. the area without liquid-was repeatedly measured (Bondza et al, 2017; Hillerdal et al, 2016; Xu et al, 2013). First, the base line recording was obtained in the four-spots setting, where each spot was read during 15 s. After 20 min, the instrument was paused and the various peptides (TM24 or SCR24) or anti-CX3CL1 antibody were added at the appropriate concentration. Then 5×10^5^ CHO_CX3CR1_ cells previously labeled with CFSE (5(6)carboxyfluorescein diacetate, succinimidyl ester, Thermofisher Scientific, Villebon-sur-Yvette, France) resuspended in 100µL of HHBSS were added in each cell dish. Then, fluorescence trace of each spot was recorded. Each experiment included control adherence experiments on L_929_ or L_CX3CL1_ cells without peptide or antibody.

### CX3CL1 binding assay using LigandTracer™

The procedure was the same as the previous one, except that the day before the experiment, 5×10^5^ CHO_CX3CR1_ cells were seeded in 35 mm Petri dishes and that the LigandTracer instrument was equipped with the 632/670 detector. When the instrument was paused, we added 100 µL containing the fluorescent CX3CL1-AF647^R^ (Almac, Craigavon, IK) and various peptides or chemokines at the appropriate concentration.

### Statistical analysis

Data were expressed as mean and standard deviations of replicates as indicated in the legend of Figs. Analysis of statistical significance were done by Student’s unpaired two-sided t-test. All statistical analysis were performed using Prism 5.2 (GraphPad Software, San Diego, USA). The levels of significance were indicated as following: *p ≤ 0.05; **p ≤ 0.005; ***p ≤ 0.0005; not significant (ns): p > 0.05.

## Acknowledgments

The authors thank Jean-Erik Guet for FRAP and FRAPP preliminary experiments, Chloé Chaudesaigues for preliminary adherence assays, Christophe Piesse for peptide synthesis, Olivier Silvie for contribution to the LigandTracer™ purchase and A. Boissonnas for valuable discussions. This work was supported by grants from the “Agence Nationale de la Recherche” (Adhekine n°09-PIRI-0003-01, CMOS n°CE-15-0019-01), from “Fondation pour la Recherche Médicale” (Equipes FRM 2016), and supports from INSERM, CNRS and UPMC.

## Author contribution

MO, PH, JJL, CC, FP and PD designed the study. PH performed gel electrophoresis. ED-M, AJ performed and analyzed the data corresponding to membrane preparations, solubilization, immuno-precipitation and native gel electrophoresis. PH and JG performed cell transfection. AC and FP preformed single-particle fluorescence experiments. MO, AC and FP performed FRAP and FRAPP experiments and analysis. MO and CH performed peptide cross-linking experiments and analysis. SIH and JJL performed CD experiments. OL performed NMR study. PF and OL performed molecular dynamics. PH, ES and NG performed adherence assays. MO, PH, AJ, JJL, CC, FP and PD analyzed data and wrote the manuscript. CC and PD collected funds.

